# Enhancing Mitochondrial Functions by Optogenetic Clustering

**DOI:** 10.1101/2022.11.22.517578

**Authors:** Kangqiang Qiu, Weiwei Zou, Zhiqi Tian, Taosheng Huang, Nien-Pei Tsai, Kai Zhang, Jiajie Diao

## Abstract

Known as the powerhouses of cells, mitochondria and its dynamics are important for their functions in cells. Herein, an optogenetic method that controlling mitochondria to form the clusters was developed. The plasmid named CRY2PHR-mCherry-Miro1TM was designed for the optogenetic system. The photoactivable protein CRY2PHR was anchored to mitochondria, via the specific organelle-targeting transmembrane domain Miro1TM. Under blue light illumination, CRY2PHR can form the oligomerization, called puncta. With the illuminated time extended, the puncta can interact, and the mitochondria were found to form clustering with reversibility and spatiotemporal controllability. The mitochondrial functions were found to enhance after the formation of optogenetic mitochondrial clusters. This method presented here provides a way to control mitochondrial clustering and raise mitochondrial functions up.

## 1. Introduction

In recent years, an emerging technique named optogenetics, based on the combination of light and genetics to control protein-protein interaction, has attracted increasing attention.[1] For the high spatial and temporal resolution of light, optogenetics possesses spatiotemporal controllability and reversibility. As the core component for optogenetics, photoactivatable proteins can initiate intracellular signal transduction by the light induced conformational change.[2] At present, optogenetics has been successfully applied to regulate a variety of cellular and organismal functions, including apoptosis, migration, differentiation, cell proliferation, gene expression, intracellular signaling, neuronal activity, and organelle transport.[3]

Mitochondria are the essential organelles for eukaryotic life.[4] They undergo fusion and fission to maintain their functions in cells.[5] Dysfunction in the mitochondrial dynamics causes organismal diseases.[6] For the importance of the mitochondrial dynamics, several methods have developed to control the mitochondrial dynamics. Genetic approaches that fragmented or elongated mitochondria by various identified proteins have been widely used for the regulations.[7] Pharmacological approaches can also regulate mitochondrial dynamics.[8] For instance, the chemical carbonyl cyanide 4-(trifluoromethoxy)phenylhydrazone (FCCP) could induce mitochondrial fission by damaging mitochondria.[9] Mitochondrial contact, also named mitochondrial clustering, a critical step for mitochondrial fusion, was reported to be obtained with a supramolecular strategy for the directed self-assembly of mitochondria by Wang group.[10] However, these methods lack the spatiotemporal controllability, and have no reversibility. Therefore, the spatiotemporal controllable and reversible method for mitochondrial dynamics’ manipulation is highly sought-after.

Recently, our group have reported that optogenetic control of mitochondria-lysosomes contacts induces mitochondrial fission, which can partially restore the mitochondrial functions of diseased cells.[3e] Inspired by such optogenetic achievement, in this work, controlling mitochondrial clustering by optogenetics was performed. Mitochondria anchored with photoactivable protein CRY2PHR were contacted under blue light illumination. The spatiotemporal controllability and reversibility of the optogenetic mitochondrial clustering were demonstrated. Interestingly, the mitochondrial functions were found to enhance after the optogenetic treatment.

## 2. Results and Discussion

### 2.1 Construct of Optogenetic System for Mitochondrial Clustering

To develop an optogenetic system for manipulating mitochondrial clustering, the blue-light-sensitive oligomerizer, cryptochrome (residues 1-498) (CRY2PHR),[2b] was opted for its no exogenous cofactors to initiate interaction. The photoactivatable CRY2PHR was anchored to the surface of mitochondria, via the specific mitochondria-targeting transmembrane domain Miro1TM (transmembrane domain of Miro1),[3a] and mCherry was used as a marker for the system’s expression, generating CRY2PHR-mCherry-Miro1TM (mCherry, a member of the mFruits family of monomeric red fluorescent proteins, Figure 1A). After transfecting different concentrations of the designed plasmid into HeLa cells, structured illumination microscopy (SIM) was used to resolve mitochondria[11]. For the appropriate fluorescent intensity and low background interference, 2.0 μg (in 200 μL DMEM, 35 mm dish; DMEM, Dulbecco’s modified eagle medium) was determined as the optimal concentration for the following transfection (Figure S1). The negligible phototoxicity of the illumination condition for the optogenetic system (60 min of continuous blue light illumination at 300 μW/cm^2^) was also verified (Figure S2). To confirm the photoactivability of CRY2PHR, the transfected cells were exposed to blue light (Figures 1B-1G, and S3). With the illumination for 5 min, the expressing proteins formed puncta gradually (Figures S3, 1C, and 1F). As the illumination time extended, the number of puncta increased (Figure S4). After 10-min exposure to blue light, the co-localization between a large punctum and a small punctum was found (Figure 1D and 1G). Incubating exposed cells in the dark for 24 h recovered the disperse state on the surface of mitochondria (Figure 1H-1J). Therefore, the results suggest that photoactivable protein CRY2PHR in the constructed optogenetic system works as design in living cells.

**Figure 1.**
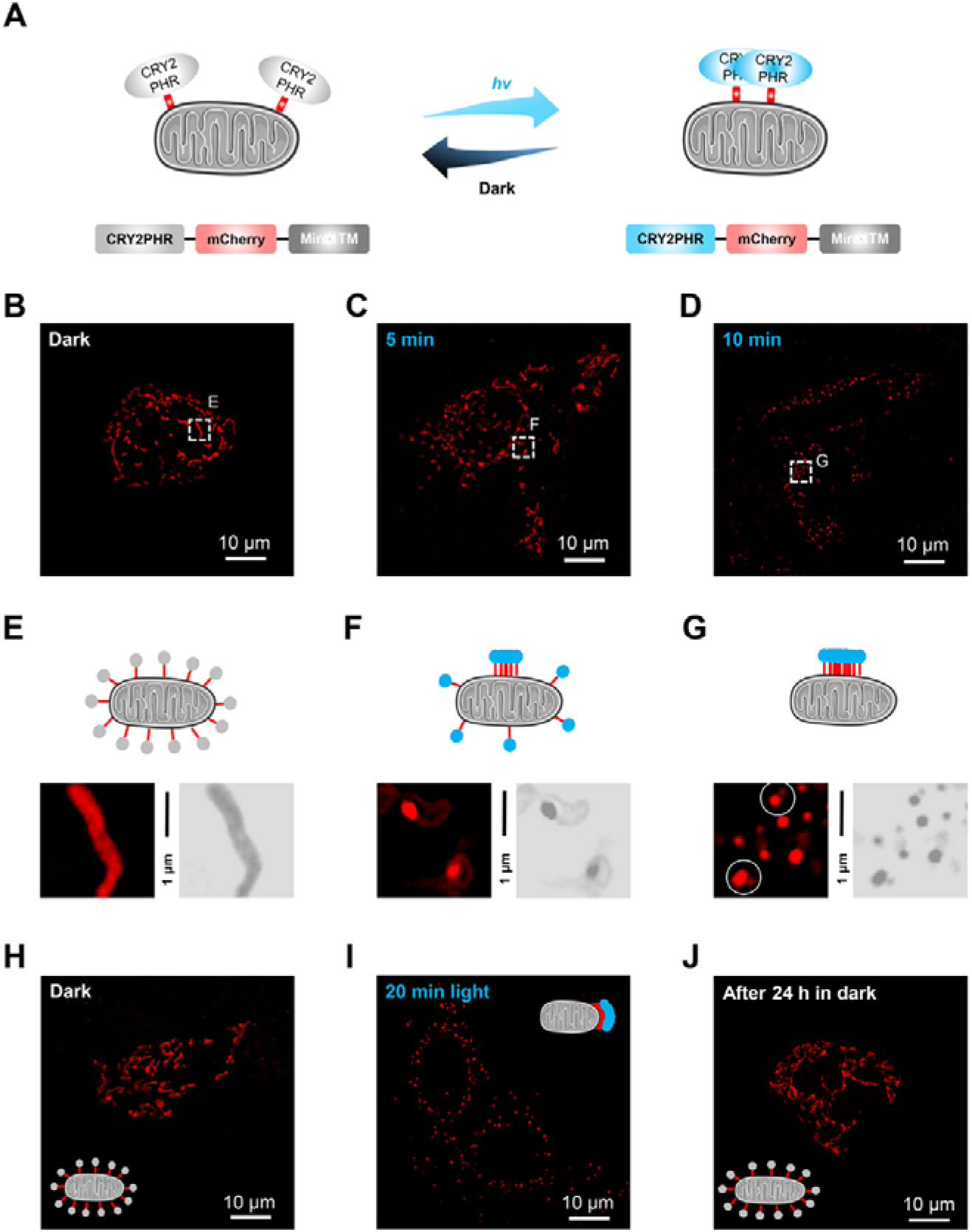
Construct of the optogenetic system for mitochondria clustering. (A) Schematic representation of the formation of CRY2PHR puncta on mitochondria under blue light exposure. Light-sensitive proteins CRY2PHR was anchored to mitochondria, via the specific organelle-targeting transmembrane domains Miro1TM. Under blue light illumination, the light-sensitive protein CRY2PHR interacted with each other to form the puncta. The fluorescent protein mCherry was used as marker for the plasmid’s expression. (B-D) Representative structured illumination microscopic images of HeLa cells expressing CRY2PHR-mCherry-Miro1TM with different time of blue light exposure at 300 μW/cm^2^; 0 min (dark) for (B); 5 min for (C); 10 min for (D). (E-G) Schematic representation of the expressing proteins on mitochondria and the partially enlarged images of Fig. 1B-1D. (H-J) The reversibility of the formation of the expressing proteins’ puncta on mitochondria. (H) Before blue-light exposure. (I) After 20 min blue-light exposure. (J) After 20 min blue-light exposure then in dark for 24 h.

### 2.2 Inducing Mitochondrial Clustering by Optogenetics

During the formation of expressing proteins’ puncta, the punctate mCherry does not depict the mitochondrial morphology. Commercial mitochondrial probes MTDR (MitoTracker™ Deep Red FM) and MTG (MitoTracker™ Green FM) were used to light up the mitochondrial dynamics. Figure 2A showed the living HeLa cells expressing CRY2PHR-mCherry-Miro1TM and stained with MTDR after blue light exposure for 20 min. There are four models for the mitochondria to contact, including single head-to-head, head-to-side, single and double side-by-side (Figure 2B). Meanwhile, the optogenetic proteins’ puncta were located at the contact sides. In the clusters, the number of mitochondria can be two, three or multiple (Figure 2C). To quantify the performance of the optogenetic system, the percentage of mitochondria in the clusters were analyzed before and after optogenetic treatment (Figures S5 and S6). As shown in Figure 2D, the percentage of mitochondria in clusters with optogenetic treatment increased almost 50%. Without light illumination, the clustering mitochondria separated, indicating that optogenetic mitochondrial clustering was reversible (Figure 2E-2H). The spatial control of the optogenetic system was demonstrated by covering half of a culture dish with tin foil paper under blue light exposure (Figure S7). To ensure that the incremental mitochondrial clustering resulted from the optogenetic manipulation, control experiments without photoactivatable proteins were performed. No noticeable changes of the percentage for mitochondria in the clusters were observed (Figure S8), demonstrating that blue light and photoactivatable proteins were essential factors for increasing mitochondrial clustering. Besides the HeLa cell line, the optogenetic mitochondrial clustering can also be performed in MCF-7 cells (Figures S9A, S10 and S11). Together, the optogenetic system efficiently induces mitochondrial clustering with reversibility and spatiotemporal controllability.

**Figure 2.**
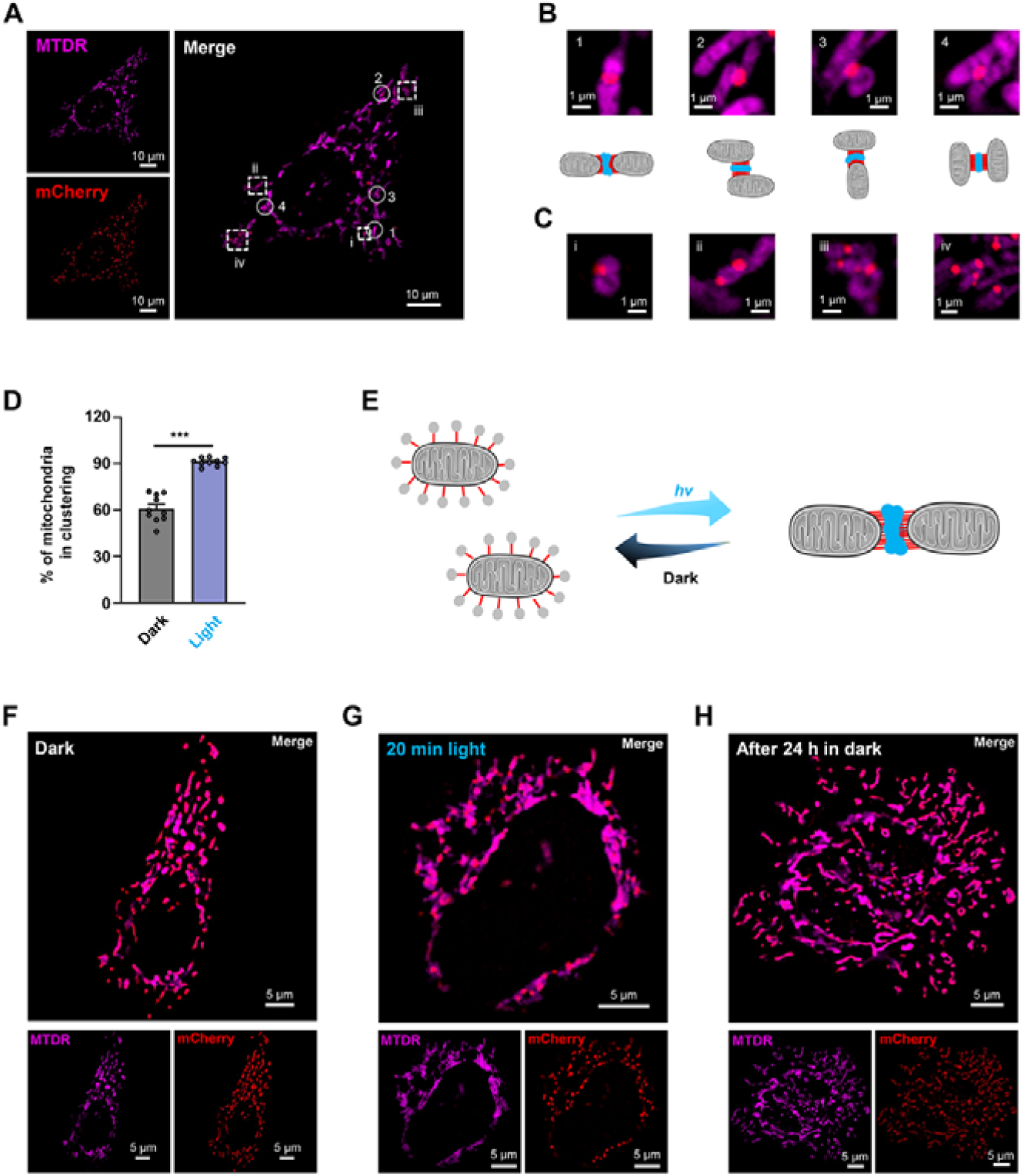
Inducing mitochondrial clustering by optogenetics. (A) Representative SIM image of living HeLa cell expressing CRY2PHR-mCherry-Miro1TM and staining MTDR. (B) Schematic representation of four contact modes in mitochondrial clusters and the partially enlarged images in Fig. 1A. (C) The partially enlarged images in Fig. 1A for the number of mitochondria to form a cluster. (D) The percentage of mitochondria in clustering for HeLa cells expressing CRY2PHR-mCherry-Miro1TM with or without blue light exposure for 20 min at 300 μW/cm^2^. Data are given as *M* ± *SEM* (*n* = 10); \*\*\**p* < 0.001 by double-tailed Student’s *t* test. (E) Schematic representation of mitochondrial clustering induced by the optogenetic system. (F-H) The reversibility of mitochondrial clustering induced by the optogenetic system. The SIM images of living HeLa cells expressing CRY2PHR-mCherry-Miro1TM and stained with MTDR. (F) Before blue-light exposure. (G) After 20 min blue-light exposure. (H) After 20 min blue-light exposure then in dark for 24 h.

### 2.3 Enhancing Mitochondrial Functions in HDFn Cells with Optogenetic Mitochondrial Clustering

It is well-known that mitochondrial dynamics can influence their functions.[5,6] MCF-7 cell line and HeLa cell line are the cancer cells. To study the mitochondrial function after optogenetic treatment, human dermal fibroblasts, neonatal (HDFn) cells were used for this study. As expected, the optogenetic system can also work in HDFn cells and the dispersive mitochondria were formed the clusters with optogenetic treatment (Figures 3A, 3B, S9B, S12 and S13).

**Figure 3.**
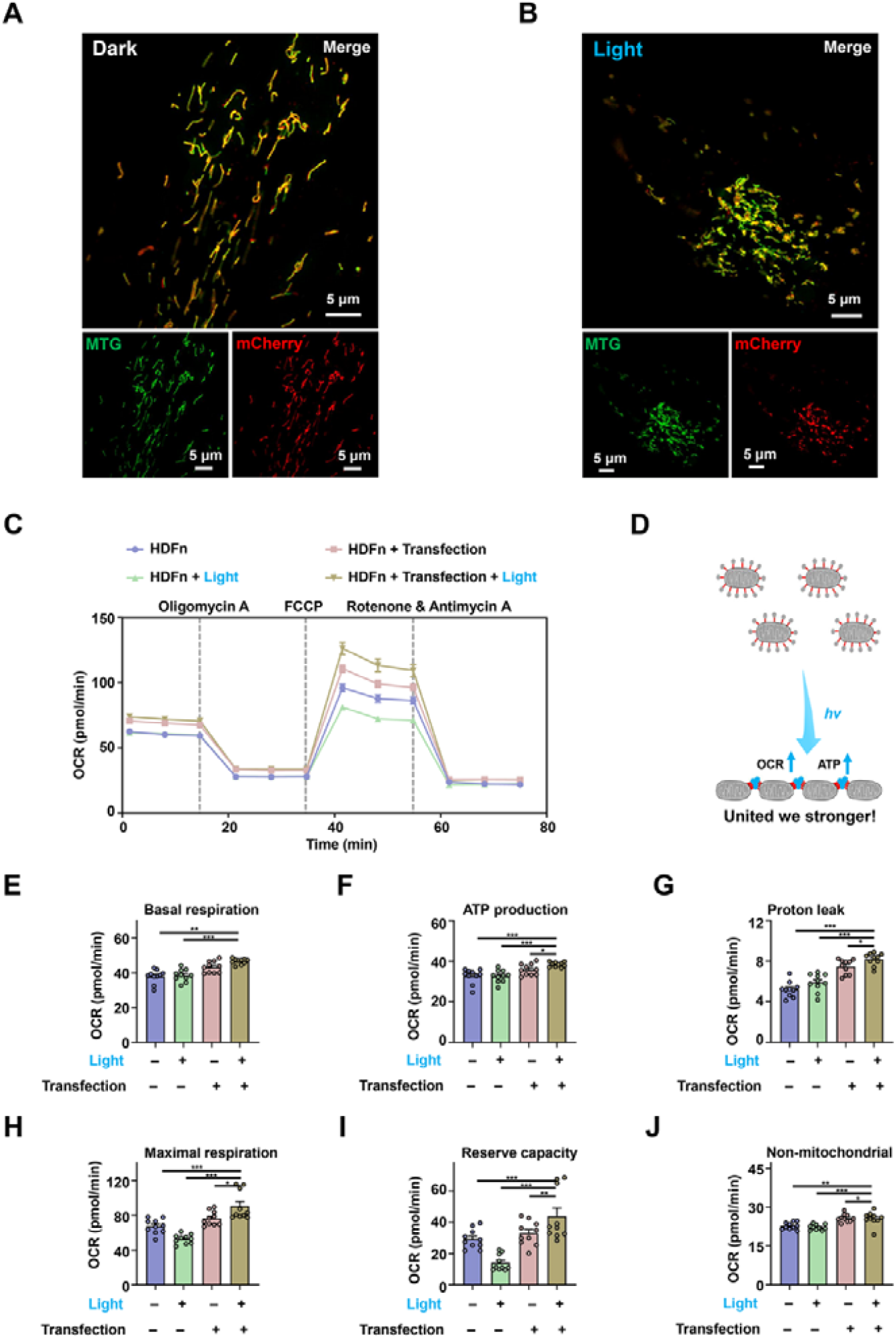
Enhancing mitochondrial functions in HDFn cells with optogenetic mitochondria clustering. (A, B) Representative SIM images of living HDFn cells expressing CRY2PHR-mCherry-Miro1TM and stained with MTG. (A) Without blue light exposure; (B) with blue light exposure at 300 μW/cm^2^ for 20 min. (C) The OCR curves of HDFn cells in real-time after different treatments. Data are given as *M* ± *SEM* (*n* = 10). OCR before the addition of oligomycin A indicates the basal respiration; OCR after the injection of FCCP denotes the maximal mitochondrial respiration capacity; and OCR after the injection of rotenone and antimycin A shows non-mitochondrial respiration. (D) Schematic representation of optogenetic treatment in HDFn cells. (E) The OCR for basal respiration of HDFn cells after different treatments. Data are given as *M* ± *SEM* (*n* = 10). (F) The OCR for ATP production of HDFn cells after different treatments. Data are given as *M* ± *SEM* (*n* = 10). (G) The OCR for proton leak of HDFn cells after different treatments. Data are given as *M* ± *SEM* (*n* = 10). (H) The OCR for maximal respiration of HDFn cells after different treatments. Data are given as *M* ± *SEM* (*n* = 10). (I) The OCR for reserve capacity of HDFn cells after different treatments. Data are given as *M* ± *SEM* (*n* = 10). (J) The OCR for non-mitochondrial respiration of HDFn cells after different treatments. Data are given as *M* ± *SEM* (*n* = 10). For (E-J), the statistical differences between the experimental groups were analyzed by double-tailed Student’s *t* test. \**p* < 0.05, \*\**p* < 0.01, and \*\*\**p* < 0.001.

The seahorse assay was employed to measure the mitochondrial oxidative phosphorylation (OXPHOS) with or without optogenetic treatment. From the oxygen consumption rate (OCR) curves in Figure 3C, there are several changes, including an initial basal OCR level, attenuated ATP-linked respiration following the addition of oligomycin (i.e., the electron transport inhibitor), a rebound of the OCR peak after addition of FCCP (i.e., the electron chain transport accelerator), and a substantial drop after a mixture of rotenone (i.e., the complex I inhibitor) and antimycin A (i.e., the complex III inhibitor) was added to discontinue the mitochondrial respiratory chain.[3e] Based on the results (Figure 3C-3J), optogenetic treatment was found to have a positive effect on the mitochondrial OXPHOS. Compared with the untreated cells, the performances in optogenetically treated cells improved 22.85% for the basal respiration (i.e., cellular energetic demand under baseline conditions, Figure 3E), 17.34% for ATP production (i.e., ATP produced to meet cellular energy needs, Figure 3F), 57.51% for proton leak (i.e., remaining basal respiration not coupled to ATP production, Figure 3G), 34.70% for maximal respiration (i.e., cells’ maximum achievable respiration rate, Figure 3H), 50.05% for Reserve capacity (i.e., the capability of the cell to respond to an energetic demand, Figure 3I), and 12.33% for non-mitochondrial respiration (i.e., oxygen consumption for a subset of cellular enzymes, Figure 3J). The results indicate that the optogenetic mitochondrial clustering system can enhance the mitochondrial functions in the normal HDFn cells.

## 3. Conclusion

In conclusion, an optogenetic method to induce mitochondrial clustering was developed in this work. With the photoactivable protein CRY2PHR on the surface, mitochondria can contact under blue light illumination to form the clusters. The percentage of mitochondria in clusters increased significantly with the optogenetic treatment. Seahorse experiment exhibited the optogenetic clustering of mitochondria can help to improve their OXPHOS. This work is a new paradigm for optogenetic manipulation of organelles to regulate cellular functions, and provides a new method for mitochondrial clustering to enhance of mitochondrial functions.

## Acknowledgements

This work was supported by the National Institutes of Health (NIH R35GM128837 to J.D.; R01GM132438 and R01MH124827 to K.Z.).

## Author Contributions

K.Q. carried out the imaging experiments. W.Z. did the Seahorse experiment. Z.T. assisted with data collection. K.Q., T.H., N-

.P.T. K.Z., and J.D. wrote the manuscript. T.H., N-.P.T. K.Z., and J.D. conceived the project. All authors participated in discussions on results and in preparing the manuscript.

