## Supporting Figures for "Enhancing Mitochondrial Functions by Optogenetic Clustering"

Table of Contents

| Experimental Section | S2-S3 |
| --- | --- |
| Supplementary Figures | S4-S16 |

Experimental Section

Materials: MitoTracker^TM^ Green FM (MTG, #M7514) and MitoTracker^TM^ Deep Red FM (MTDR, #M22426) were purchased from Invitrogen (Thermo Fisher Scientific, USA). Penicillin–streptomycin (#15140163, 10,000 units/mL), fetal bovine serum (FBS, #26140079), and DMEM (#11965092) were all purchased from Gibco (Thermo Fisher Scientific, USA). Phosphate-buffered saline (PBS, #SH30256.01) was purchased from Hyclone (GE Healthcare Life Sciences, USA). CRY2PHR-mCherry-Miro1TM (Addgene plasmid, #102247) was a gift from Bianxiao Cui. TurboFect^TM^ Transfection Reagent (#R0532) was purchased from Thermo Scientific (Thermo Fisher Scientific, USA).

Cell Culture and Transfection: The HeLa cell line (a gift from Dr. Carolyn M. Price, University of Cincinnati), MCF-7 cell line (a gift from Prof. Jun-Lin Guan, University of Cincinnati) and HDFn cell line were cultured in DMEM containing 10% FBS and 100 units/mL of penicillin–streptomycin in a 5% CO_2_ cell incubator (Thermo Fisher Scientific, USA) with 100% humidity at 37 °C. Transfection was performed using TurboFect^TM^ Transfection Reagent according to the manufacturer's protocol. CRY2PHR-mCherry-Miro1TM was mixed with 6 μL TurboFect in 200 μL DMEM medium (without FBS) for 20 min. The DNA/Turbofect mixtures were added to the cell cultures drop-wise and incubated for 6 h before replenishment with complete culture medium.

Structured Illumination Microscopy Imaging: All SIM images were performed with a Nikon structured illumination microscopy (N-SIM, version AR5.11.00 64bit, Tokyo, Japan), a 3D-SIM equipped with an Apochromat 100×/1.49 numerical aperture oil-immersion objective lens and solid-state lasers (488 nm, 561 nm, 640 nm, the output powers at the fiber end: 15 mW). Images were captured using Nikon NIS-Elements 512 × 512 using Z-stacks with a step size of 0.2 μm and the raw images were reconstructed and processed with NIS-Elements AR Analysis (version AR5.11.00 64bit). The green channel images with emission bandwidth at 500-550 nm were excited by a 488 nm laser for MTG. The red channel images with emission bandwidth at 570-640 nm were excited by a 561 nm laser for mCherry. The deep red channel images with emission bandwidth at 660-735 nm were excited by a 640 nm laser for MTDR. Cells were seeded on glass-bottomed culture dishes (MatTek; P35G-1.5-14-C) for 24 h to adhere. Staining with commercial dyes were performed for 30 min. Before imaging, cells were washed with PBS 3 times. For illumination, a homemade blue light emitting diode (LED) array was used with 300 μW/cm^2^. The imaging data analysis was performed with ImageJ.

Oxygen Consumption Rate Assay: To examine the mitochondrial OXPHOS function under blue light exposure, the oxygen consumption rate (OCR) of HDFn cells expressing CRY2PHR-mCherry-Miro1TM was measured using a Seahorse XFe-96 Analyzer (Agilent Technologies, USA) in real time. HDFn cells (two groups, one for illumination and one for dark) and HDFn cells expressing CRY2PHR-mCherry-Miro1TM (two groups, one for illumination and one for dark) were seeded in a XFe-96 cell culture plate (Agilent Technologies, USA) at 0.8 × 10^4^ cells/well in DMEM with 10% FBS. After 48 h, the medium was removed and replaced with the warmed unbuffered "Seahorse medium" (XF DMEM with 1 mM sodium pyruvate, 10 mM glucose and 2 mM L-glutamine) at pH 7.4. Before the measurement, one group of HDFn cells and one group of HDFn cells expressing plasmid were exposed to blue light (300 μW/cm^2^) for 20 min. Then the OCRs of the cells were assessed using the XF Cell Mito Stress Test Kit (Agilent Technologies, USA) according to the manufacturer's instruction. Where indicated, the cells were treated with 1 μM oligomycin A (the electron transport inhibitor), 2 μM FCCP (the electron chain transport (ETC) accelerator), and 500 nM rotenone (the complex I inhibitor) with 1 μM antimycin A (the complex III inhibitor, all are from Agilent Technologies).

Cell Viability Test: Cell Counting Kit-8 (CCK-8, Dojindo Molecular Technologies, Inc., Japan) was used to determine the photocytotoxicity of blue-light. The cells (HeLa cells, MCF-7 cells and HDFn cells) were seeded into two groups in a 96-well plate with 1 × 10^4^ cells/well. After 24 h to adhere, the light group was exposed to blue-light (300 μW/cm^2^) for 60 min, while the dark group was covered with tin foil sheet and left unchanged. Then 10 μL of CCK-8 solution was added to each well, the culture plate was incubated for 1 h. Absorbance at 490 nm was determined with the Synergy Mx microplate reader (BioTek Instruments, Inc., USA).

Data Analysis: Statistical significance of data was evaluated using Student's *t* test. Data were presented as *M* ± *SEM*. Statistics and graphing were performed using Prism 8 (GraphPad) or Excel (Microsoft). The images were analyzed by ImageJ-win64 (NIH Image). All images were assembled using PowerPoint 2016 software (Microsoft).

Supplementary Figures

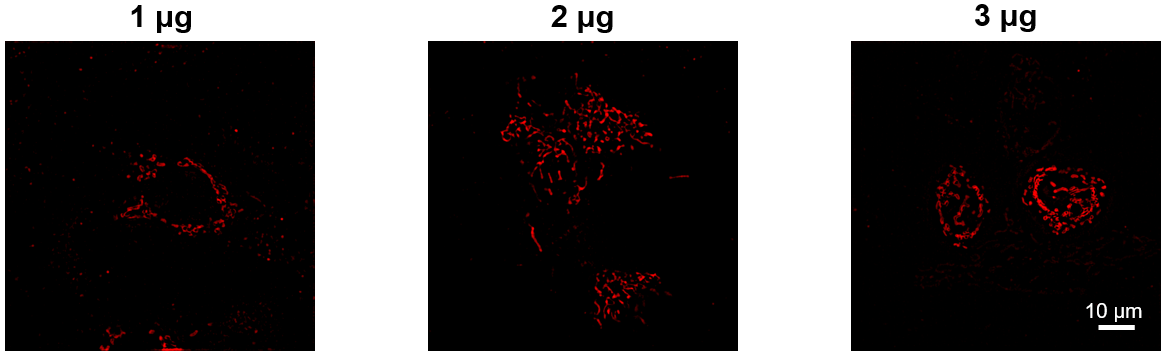

**Figure S1.** The SIM images of living HeLa cells expressing different concentrations of CRY2PHR-mCherry-Miro1TM. All images shared the same scale bar.

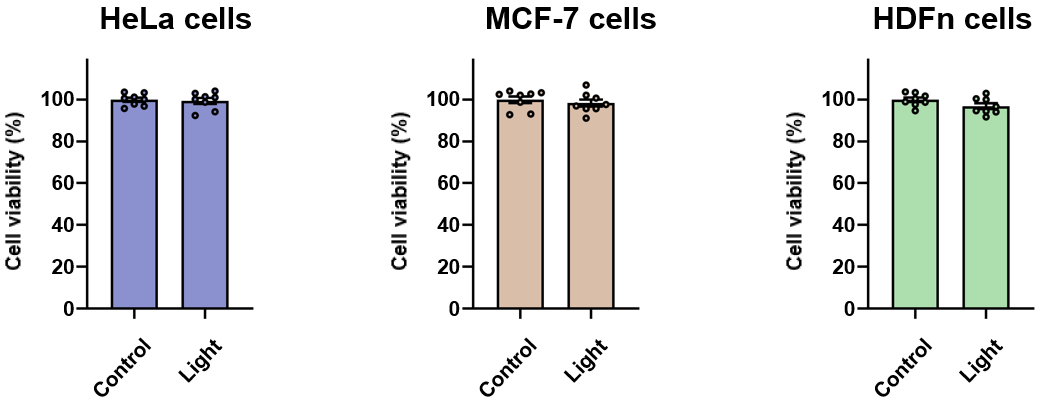

**Figure S2.** The viability of HeLa cells, MCF-7 cells, and HDFn cells with/without blue-light exposure for 60 min. Data are presented as *M* ± *SEM*.

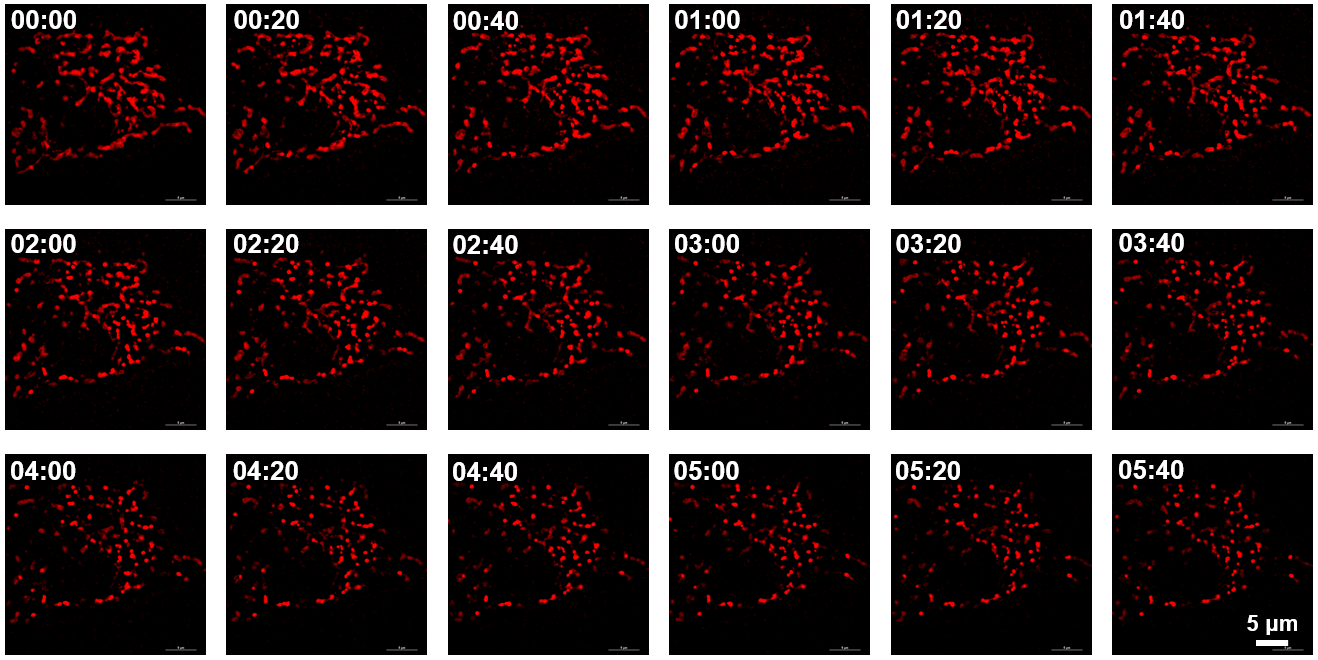

**Figure S3.** The time-lapse SIM images of a same HeLa cell expressing CRY2PHR-mCherry-Miro1TM under blue light exposure. All images shared the same scale bar.

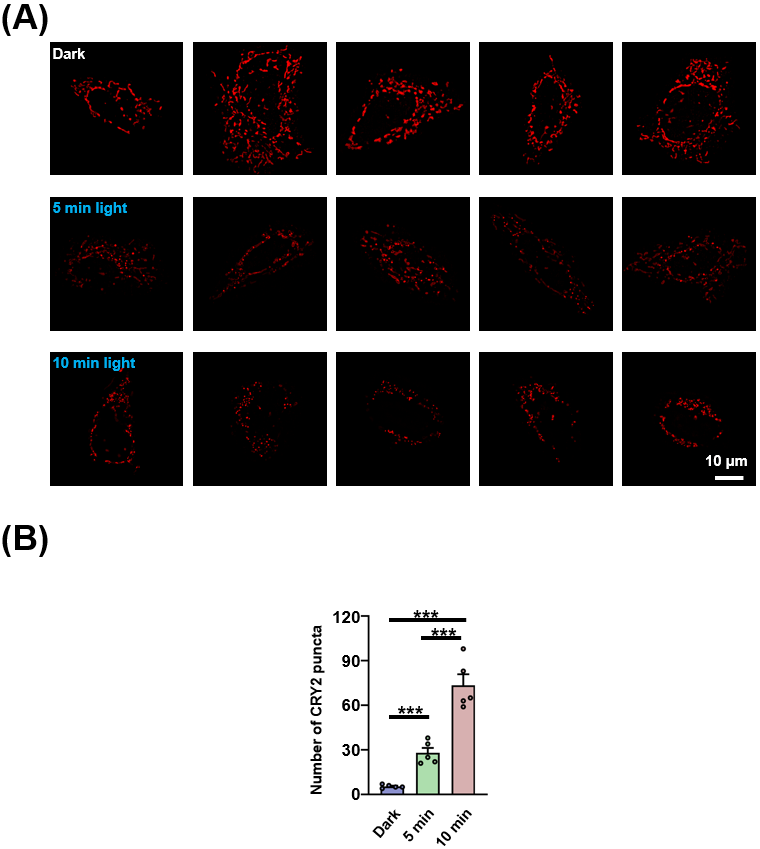

**Figure S4.** The increasing number of the expressing proteins' puncta under blue light exposure. (A) The SIM images of living HeLa cells expressing CRY2PHR-mCherry-Miro1TM with blue light exposure for different time. (B) Quantitative analysis of the number of puncta in Figure S4A.The statistical differences between the experimental groups were analyzed by double-tailed Student’s *t* test. ****p* < 0.001.

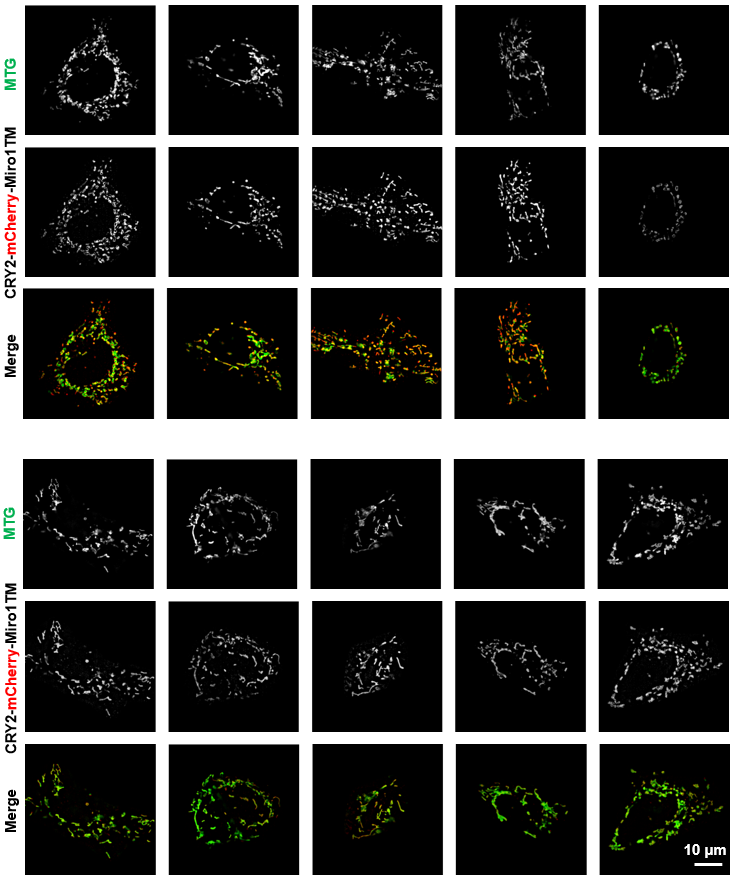

**Figure S5.** Data set for Dark group in Figure 2D. The SIM images of living HeLa cells expressing CRY2PHR-mCherry-Miro1TM and stained with MTG.

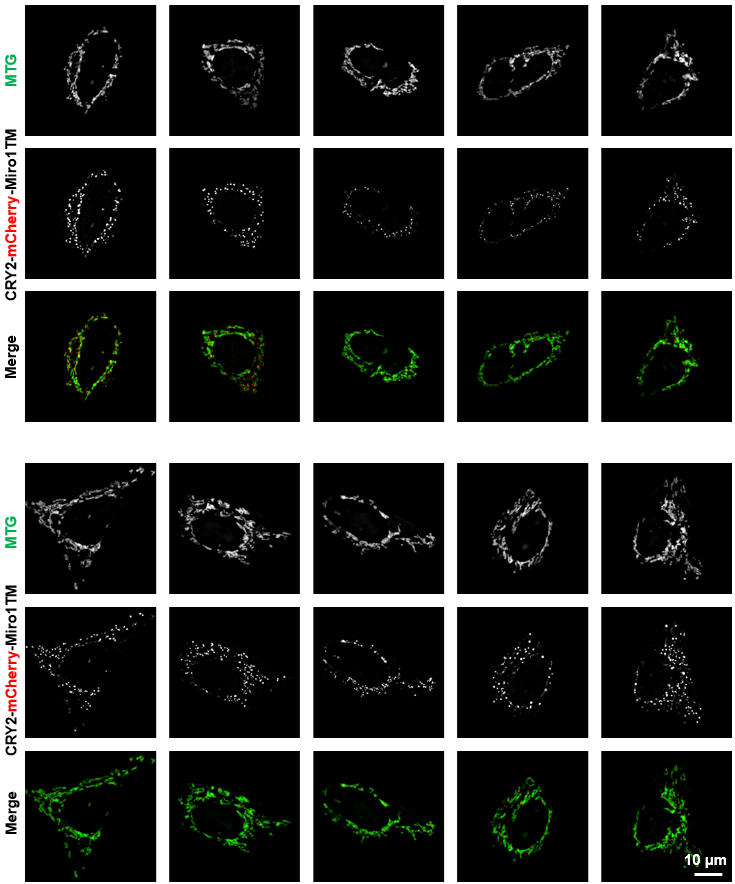

**Figure S6.** Data set for Light group in Figure 2D. The SIM images of living HeLa cells expressing CRY2PHR-mCherry-Miro1TM and stained with MTG under blue light for 20 min.

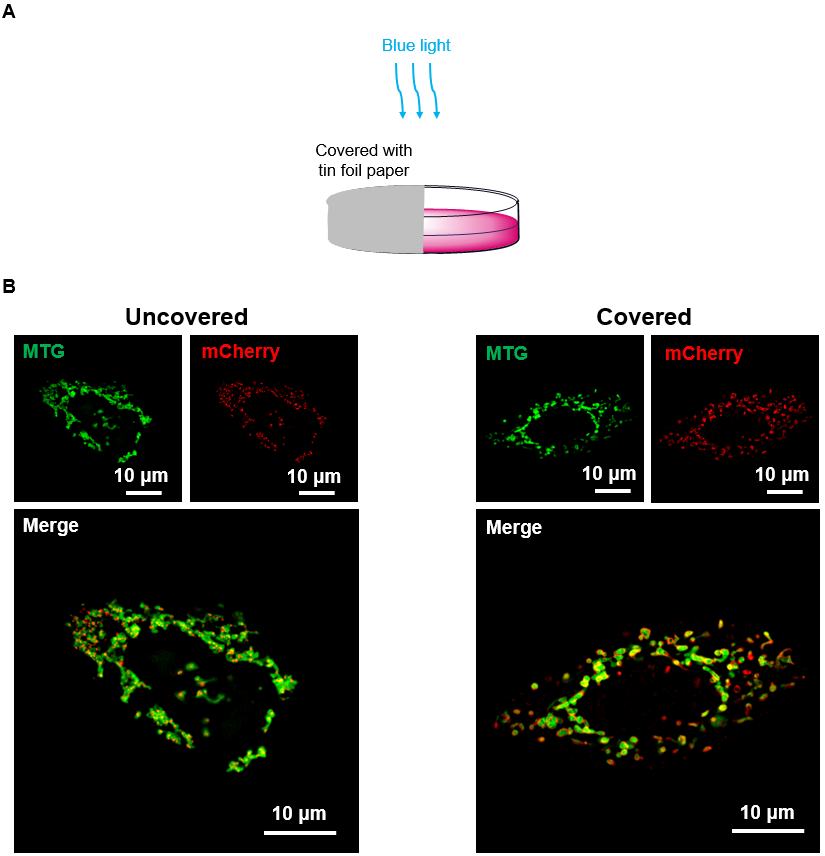

**Figure S7.** (A) Schematic representation of the spatial control experiment, where half of the culture dish was covered with tin foil paper to darken that portion of the dish. (B) The SIM images of HeLa cells expressing CRY2PHR-mCherry-Miro1TM and stained with MTG by spatial control with or without light exposure for 20 min.

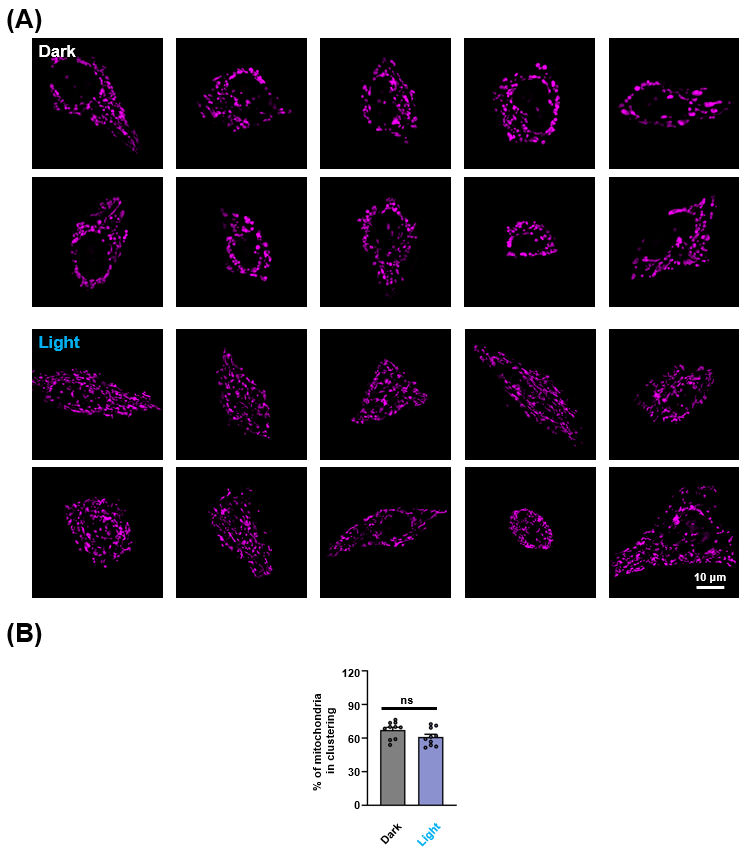

**Figure S8.** (A) The SIM images of HeLa cells stained with MTDR with or without blue light exposure for 20 min. (B) Quantitative analysis of the percentage of mitochondria in Fig. S8A. Data are given as *M* ± *SEM* (*n* = 10); No significant by double-tailed Student’s *t* test.

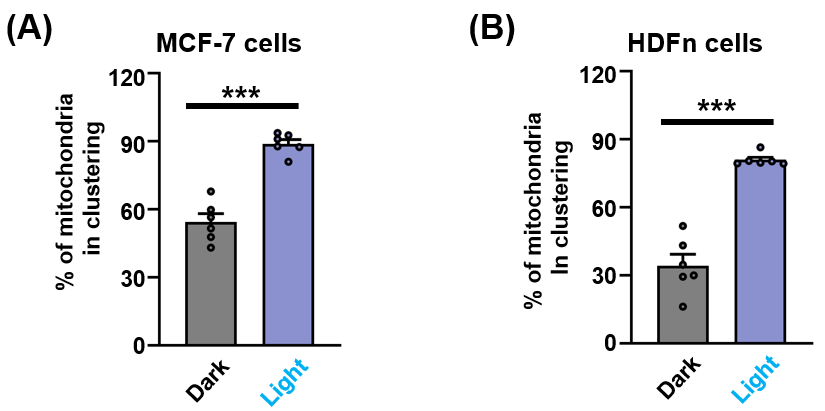

**Figure S9.** Quantitative analysis of the percentage of mitochondria in clusters for MCF-7 cells or HDFn cells expressing CRY2PHR-mCherry-Miro1TM and staining MTG with or without blue light exposure. Data are given as *M* ± *SEM* (*n* = 6); ****p* < 0.001 by double-tailed Student’s *t* test.

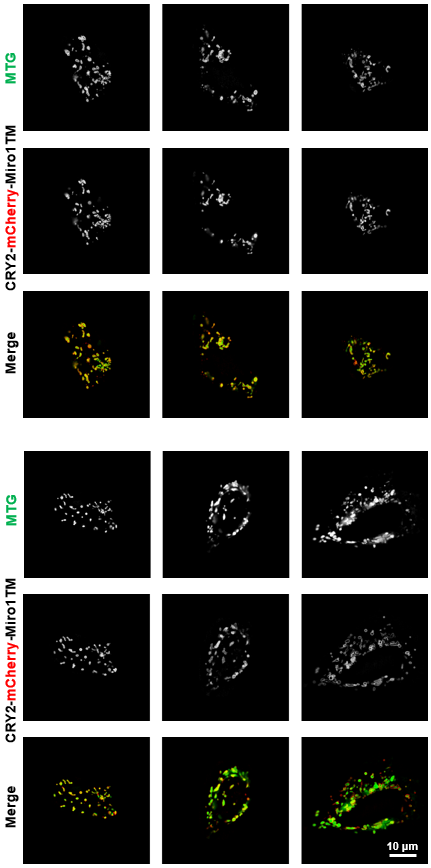

**Figure S10.** Data set for Dark group in Figure S9A. The SIM images of living MCF-7 cells expressing CRY2PHR-mCherry-Miro1TM and stained with MTG.

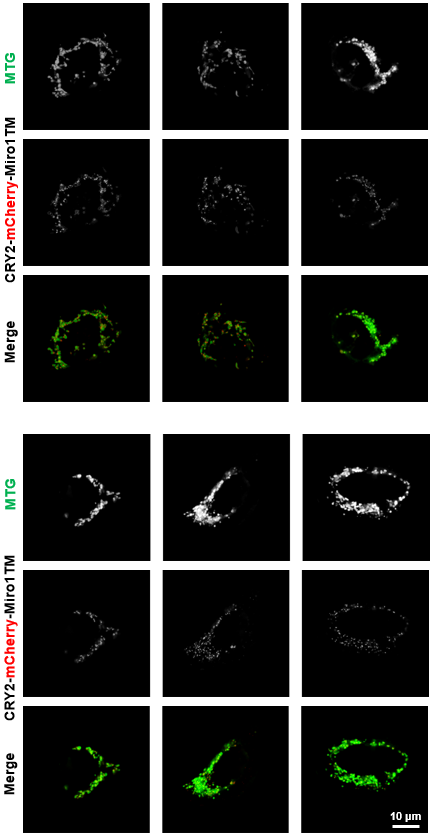

**Figure S11.** Data set for Light group in Figure S9A. The SIM images of living MCF-7 cells expressing CRY2PHR-mCherry-Miro1TM and stained with MTG under blue light for 20 min.

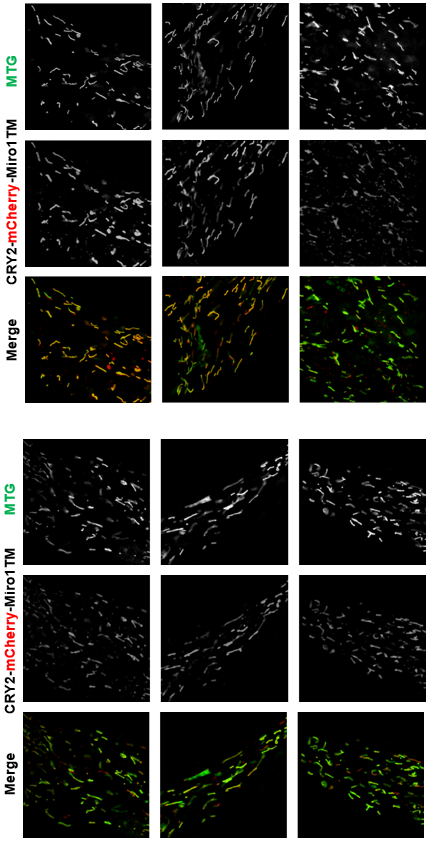

**Figure S12.** Data set for Dark group in Figure S9B. The SIM images of living HDFn cells expressing CRY2PHR-mCherry-Miro1TM and stained with MTG.

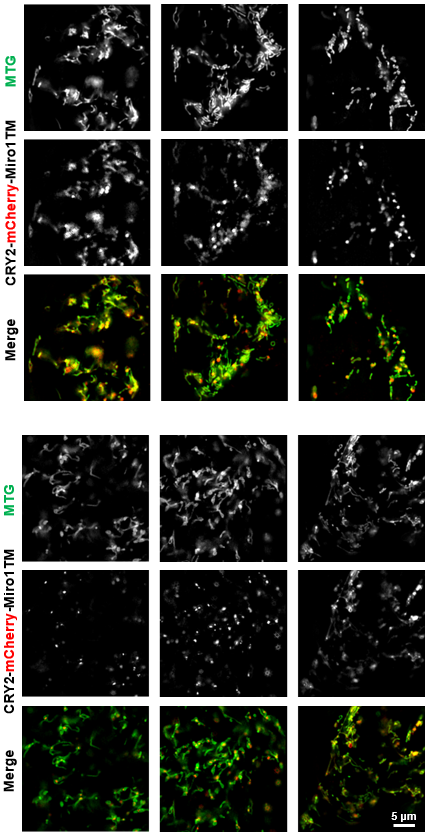

**Figure S13.** Data set for Light group in Figure S9B. The SIM images of living HDFn cells expressing CRY2PHR-mCherry-Miro1TM and stained with MTG under blue light for 20 min.
